# Thermal acclimation to warmer temperatures can protect host populations from both further heat stress and the potential invasion of pathogens

**DOI:** 10.1101/2022.05.04.488533

**Authors:** Tobias E. Hector, Marta S. Shocket, Carla M. Sgrò, Matthew D. Hall

## Abstract

Phenotypic plasticity in response to shifts in temperature, known as thermal acclimation, is an essential component of the ability of a species to cope with environmental change. Not only does this process potentially improve an individual’s thermal tolerance, it will also act simultaneously on various fitness related traits that determine whether a population increases or decreases in size. In light of global change, thermal acclimation therefore has consequences for population persistence that extend beyond simply coping with heat stress. This particularly important when we consider the additional threat of parasitism associated with global change, as the ability of a pathogen to invade a host population depends on both its capacity to proliferate within a host and spread between hosts, and thus the supply of new susceptible hosts in a population. Here, we use the host *Daphnia magna* and its bacterial pathogen *Pasteuria ramosa* to investigate how thermal acclimation may impact various aspects of host and pathogen performance at the scale of both an individual and the population. We independently test the effect of maternal thermal acclimation and direct thermal acclimation on host thermal tolerance, measured as knockdown times, as well as host fecundity and lifespan, and pathogen infection success and spore production. We find that direct thermal acclimation enhances host thermal tolerance and intrinsic rates of population growth, despite a decline observed for host fecundity and lifespan. Pathogens, on the other hand, faired consistently worse at warmer temperatures at the within-host scale, and also in their potential to invade a host population. Our results suggest that hosts could benefit more from warming than their pathogens, but highlight that considering both within- and between-host thermal performance, including thermal tolerance and fitness traits, is needed to fully appreciate how increasing thermal variability will impact host and pathogen populations.

## Introduction

Processes at every level of biological organisation are fundamentally shaped by temperature, from the rate at which physiological process occur within the body, to the growth and persistence of a population or community (Angilletta 2009; Chown et al. 2010; Somero 2010; Colinet et al. 2015; Sinclair et al. 2016; Vázquez et al. 2017). This is particularly true for host-pathogen interactions. A host or pathogen’s response to changing temperatures depends both on metrics of individual performance such as host fecundity, lifespan, and pathogen proliferation, as well as population level processes such as host population growth rate or the capacity of a pathogen to spread between hosts (Anderson and May 1986; Mideo et al. 2008; Hall and Mideo 2018). Yet, studies often only look at snapshots of these traits, or focus on the individual scale (but see Cuco et al. 2018; Agha et al. 2018; Shocket et al. 2018a; 2019), such as how temperature impacts host fecundity or survival, or how temperature alters pathogen proliferation or virulence (Elliot et al. 2002; Mitchell et al. 2005; Laine 2007; Vale et al. 2008; Vale and Little 2009; Hector et al. 2019). Viewing each component in isolation has the potential to be misleading. While population level processes intrinsically depend on individual responses to temperature, one does not necessarily predict the other (Mideo et al. 2008; Wolinska and King 2009; Penczykowski et al. 2016; Hall and Mideo 2018).

For a host, exposure to rising non-lethal temperatures, through the process of acclimation, allows individuals to shift their thermal optima or maxima, potentially acting as a buffer against future thermal stress (Sinclair et al. 2016; Sgrò et al. 2016; Rohr et al. 2018). However, thermal acclimation to warmer temperatures will also typically accelerate the pace of life, leading to earlier reproductive output and shortened lifespans (Zwaan et al. 1992; Angilletta et al. 2004; Stoks et al. 2014), which potentially changes patterns of population growth. Similar processes are also likely to occur for a pathogen. Exposure to higher temperatures can increase the rates at which pathogen can encounter and infect a host (Shocket et al. 2018a,b), but will also accelerate the infection process, by increasing pathogen replication rates while at the same time increasing the virulence of a pathogen and shortening the duration of infection (Mitchell et al. 2005; Fels and Kaltz 2006; Vale et al. 2008; Cuco et al. 2018). It is now clear that pathogen exposure can also severely reduce a host’s capacity to cope with thermal stress (Greenspan et al. 2017; Hector et al. 2019), meaning that population persistence will not only depend on a host’s thermal performance in isolation, but also on the simultaneous impact of disease exposure (Gehman et al. 2018; Hector et al. 2019; 2020).

The response of hosts and pathogens to increasing temperatures will depend on a balance between the effects of thermal acclimation on their thermal tolerance and thermal performance, versus effects on traits that underlie whether a host or pathogen population will increase or decrease as temperatures change. When assessing the benefits or costs of thermal acclimation, therefore, the potential duality of host and pathogen performance at the individual and population levels needs to be considered. This can be achieved by comparing changes in thermal tolerances with host and pathogen vital metrics. For hosts, life tables can be used to calculate population growth and death rates in order to evaluate how temperature exposure may influence host population dynamics (McCallum 2000; Civitello et al. 2013; Shocket et al. 2018a). In turn for a pathogen, estimates of infection success and proliferation can be integrated via an epidemiological model into the basic reproduction number, *R_0_*, which captures the potential of a pathogen to spread through a completely susceptible host population (Anderson and May 1986). Rarely, however, has the impact of thermal acclimation on thermal stress resistance been considered in unison with metrics of how well a host or pathogen population might perform (Klockmann et al. 2017; Cavieres et al. 2020). Evidence for whether thermal acclimation improves or impairs thermal stress resistance when a host is simultaneously exposed to disease is likewise limited (see Greenspan et al. 2017).

Thermal acclimation is also not restricted to the direct effects of temperature on a host or pathogen during infection. There is now growing evidence that both trans-generational and developmental temperature exposure can lead to shifts in both thermal performance and many characteristics of infection (Hoffmann et al. 2012; van Heerwaarden et al. 2016; Sgrò et al. 2016; Beaman et al. 2016; Kellermann et al. 2017; Moghadam et al. 2019). In *Drosophila melanogaster*, for example, developmental temperature exposure has been shown to have a greater influence on adult thermal tolerance than direct thermal acclimation (Slotsbo et al. 2016; Kellermann et al. 2017). Thermal acclimation can also impact disease traits across generations. For example, host resistance to infection can be mediated by the thermal environment experienced in the maternal generation (Garbutt et al. 2014). Whilst, for a pathogen, the temperature experienced during one infection cycle can lead to changes in the infectivity of the spores involved in subsequent cycles of infection, analogous to trans-generational effects (Altman et al. 2016; Shocket et al. 2018b). Maternal and developmental acclimation, therefore, has the potential to be an important mechanism for preparing populations to cope with future environmental conditions (Sgrò et al. 2016). However, it is unclear how thermal acclimation prior to infection will influence the impact of infection on host resistance to thermal stress, and, in addition, affect host and pathogen performance across individual to population level scales.

In this study, we contrasted how thermal acclimation, before and during infection, shapes host thermal tolerance and host and pathogen individual performance, versus the potential for growth of host and pathogen populations. To address these questions, we used the water flea *Daphnia magna* and its bacterial pathogen *Pasteuria ramosa. Daphnia* have been shown to be able to mount strong plastic thermal acclimation responses in their upper thermal limits (Williams et al. 2012; Yampolsky et al. 2014; Burton et al. 2020), and temperature is known to mediate their response to infection (Mitchell et al. 2005; Vale et al. 2008; Allen and Little 2011). *Pasteuria ramosa* is a natural bacterial pathogen of *Daphnia*, which has distinct genotypes known to vary in various aspects of within-host performance, including infection rates and spore production (Clerc et al. 2015; Hall and Mideo 2018), both of which have been shown to be sensitive to temperature stress (Vale et al. 2008; Vale and Little 2009; Hector et al. 2019). We used two levels of acclimation, the first being a maternal and developmental acclimation treatment prior to infection, and the second being direct thermal acclimation on focal individuals including over the infection period. These two acclimation levels allowed us to test the separate effects of maternal and developmental acclimation prior to infection, and direct thermal acclimation during infection, on host thermal tolerance alongside other important components of host and pathogen thermal performance.

We first measured thermal tolerance of infected and uninfected individuals as knockdown time under heat shock for each thermal acclimation regime. Next, we measured host lifespan and fecundity for infected and uninfected individuals, as well as within-host pathogen spore loads and infection success for each thermal acclimation treatment. For hosts, we then used this life table data of lifespan and fecundity to calculate population growth rates and death rates under each thermal acclimation treatment to evaluate how temperature exposure may influence host population dynamics (McCallum 2000). For the pathogen, we incorporated estimates of infection success and proliferation into an epidemiological model to estimate a metric for the potential for disease spread through a host population (i.e., Anderson and May 1986), the basic reproduction number (*R_0_*), under each thermal acclimation treatment. The ability of a pathogen to persist in a host population is influenced by both host population dynamics and pathogen fitness, and therefore depends on its ability to proliferate within a host, its capacity to spread between hosts, and the supply of new susceptible hosts in a population. Together, these measures allow us to contrast how thermal acclimation can impact various metrics of host and pathogen thermal performance and fitness at an individual scale, and how they combine to determine the population scale outcomes.

## Methods

### Host and pathogen

The cyclically parthenogenic crustacean *Daphnia magna* Straus is commonly found in both fresh and brackish waters, including shallow pools and large lakes, across Eurasia (Ebert 2005). *Pasteuria ramosa* Metchnikoff is a Gram-positive bacterial pathogen of *D. magna* that enters the host during filter feeding before reducing lifespan and fecundity of its host (Hall and Ebert 2012; Clerc et al. 2015; Ebert et al. 2016). At host death millions of spores are released into the environment where exclusively horizontal transmission takes place, which itself depends on the interplay between the pathogen’s ability to produce mature transmission spores and its virulence (Hall and Mideo 2018). In this study we used *Daphnia* genotype BE-OMZ-M10 infected with one of three *P. ramosa* genotypes (C1, C14 and C20). These pathogen genotypes were chosen because they display a range of virulence and transmission potentials (Clerc et al. 2015; Hall and Mideo 2018) and also vary in the extent to which they reduce host thermal tolerances (Hector et al. 2019).

Before the experiments, female *Daphnia* taken from stock culture were placed individually in 70-mL jars filled with 50 mL of Artificial *Daphnia* Medium (ADaM; Ebert et al. 1998) for three generations to minimise trans-generational effects. *Daphnia* were changed into fresh ADaM twice a week and fed with algae (*Scenedesmus sp*.) daily. Food levels were increased from one million cells at birth to eight million by age 14 days to meet the growing energy needs of the animals. *Daphnia* were maintained under standard conditions (20°C, 16L:8D) and repositioned within the incubator regularly in order to minimise any positional effects.

### Experimental animals, thermal acclimation, and infection

Thermal acclimation began in the maternal generation. On the day of birth, F0 (maternal generation) individuals were taken from clutches 3–5 of the standardised animals and maintained at either 20°C or 25°C (maternal/developmental temperature treatment, hereafter maternal acclimation). Experimental F1 animals were then collected from clutches 3–5 of the acclimated mothers on the day of birth and placed at either 20°C or 25°C (focal acclimation temperature) in a fully factorial design, resulting in four thermal acclimation treatments (20-20, 20-25, 25-20 & 25-25°C). The outcome of the thermal acclimation treatments was that the first temperature experienced would involve maternal and developmental effects (because *Daphnia* develop within the mother and are live born), whilst the second temperature experienced would result in direct thermal acclimation on the focal animals which lasted the whole of their life, including over the infection period. Experimental animals were kept at their acclimation temperatures from birth until either being used in thermal tolerance assays or until death (see below for details).

A total of 1008 females were set up in the experimental generation in a fully factorial design, with 63 individuals per treatment (2 maternal temperatures x 2 focal temperatures x [3 pathogens + uninfected controls]). Individual *Daphnia* were infected with 40,000 *P. ramosa* spores over two days (20,000 per day) starting three days after birth. Infection took place in 70-mL jars filled with 20 mL ADaM for three days, after which all animals were transferred to fresh ADaM and maintained as described above.

### Thermal tolerance assays

Static heat shock was used to measure thermal tolerance as heat knockdown time of *Daphnia* from all treatments described above. Individual *Daphnia* were placed in 5-mL glass fly vials covered in mesh and immersed in a constantly agitated water bath filled with ADaM and set to 37°C (Hector et al. 2019). All individuals were monitored constantly throughout the assay and time until knockdown, starting from when they were first placed in the water bath, was recorded when there was no visible movement from the *Daphnia* (Yampolsky et al. 2014; Hector et al. 2019). A total of 36 *Daphnia* per treatment were chosen at random to measure thermal tolerance. Three individuals per treatment could be measured per assay run, so 12 assay runs were conducted over three consecutive days. All animals were between 19 and 21 days post-infection at the time of the assays.

### Host and pathogen disease traits

The remaining *Daphnia* that were not used in the thermal tolerance assays were kept at their respective focal acclimation temperatures until death. From birth, all animals were checked daily for deaths, and any dead animals were frozen in 500 μL of RO water for later bacterial spore counting. Twice-weekly all individuals also had their offspring counted. This gave us four important metrics of host and pathogen fitness for each temperature by pathogen treatment combination: host lifespan, host age-specific fecundity, pathogen spore loads at host death, and infection rates.

### Bacterial spore counts

Bacterial spore counts were quantified using an Accuri C6 Flow Cytometer (BD Biosciences, San Jose, California). Infected animals were thawed and homogenised in 500 μL of RO water. Then, 10 μL of this sample was pipetted into 190 μL of 5mM EDTA in a 96-well plate. For each run, 6 samples were counted with every fourth well containing only EDTA as a wash step. A combination of gates based on fluorescence (via the 670 LP filter) and side scatter (cell granularity) were used to identify mature spores based on their distinct size, morphology, and fluorescence, compared to immature spores, algae or animal debris. Each sample was counted twice and counts were averaged, and then used to calculate total spore load per infected individual. Samples were also checked under a microscope to determine whether individuals contained mature transmission spores, which would count as a successful infection, or only contained undeveloped spores that would be unable to infect another host and therefore represents an unsuccessful infection.

### Host population growth and between host disease spread: the model and the parameters

We investigated how thermal acclimation would impact disease spread through a population using a model. This model tracks changes in the density of susceptible (*S*) and infected (*I*) hosts and environmental pathogen spores (*Z*) (Hall et al. 2009; Civitello et al. 2013)

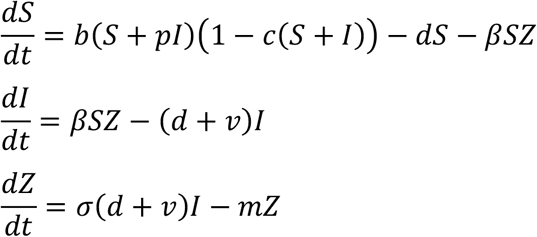

where susceptible hosts increase in a density dependent manner, which itself is dependent on a maximum birth rate, *b*, and the strength of density dependence, *c*. Infection leads to a reduction in fecundity (0 ≤ *p* ≤ 1). Susceptible hosts die at a constant background rate, *d*. Susceptible hosts become infected dependent on an infection rate β, and contact with spores, *Z*. Infected hosts die at a constant background rate (*d*) in addition to the virulence of infection (*v*). Spores are released into the environment from dead hosts with a spore load σ, and are lost from the environment (due to degradation) at rate, *m*.

From this model we can calculate a metric that informs us about a pathogens potential to spread through an entirely susceptible population, otherwise known as a reproductive ratio, *R_0_* (Anderson and May 1986). Larger values of *R_0_* suggest the potential for larger epidemics, which will have greater impacts on host and pathogen populations (Anderson and May 1986). We use *R_0_* as a qualitative indicator of how our thermal acclimation treatments would impact various between-host processes of host and pathogen populations, and their relative contribution to the potential for disease spread. From our model above, *R_0_* is

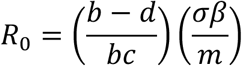

which is dependent on susceptible host density in the absence of disease, (*b* – *d*)/(*bc*), and three epidemiological traits, (σβ/*m*). *R_0_* will increase if there are increases in host birth rate, *b*, pathogen transmission rate, β, or pathogen spore loads, σ. *R_0_* decreases if there are increases in host death rate, *d*, the rate of pathogen loss from the environment, *m*, or the strength of density dependence on host birth rate, *c* (Civitello et al. 2013).

### Model parameterization

To calculate *R_0_* for each thermal acclimation treatment and pathogen genotype we calculated various important parameters from the model above using data from animals that we observed from birth till death. First, we calculated the intrinsic rate of increase (population growth rate – *r*) for each unexposed (susceptible) host individual in each temperature treatment by solving the Euler-Lotka equation

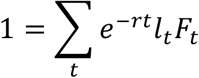

where *l_t_* is the proportion of individuals in a cohort surviving to day *t*, and *F_t_* is the average fecundity at day *t* for each treatment group. Following the methodology of Shocket *et al*. (Shocket et al. 2018a) we used a simplified version of this equation to calculate the intrinsic rate of increase for each individual (rather than for each whole population), where for each single individual *l_t_* always equals 1 while the animals remains alive and *F_t_* is the fecundity of each individual at day *t*, yielding the equation

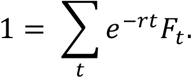

We next calculated instantaneous death rate (*d*) for our uninfected (susceptible) hosts in each temperature treatment assuming time until death followed an exponential distribution, where the likelihood of a constant death rate (*d*) is calculated from our time until death (lifespan) data under each temperature treatment (Civitello et al. 2013; Shocket et al. 2018a)

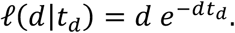

Birth rate (*b*) for uninfected (susceptible) hosts was then calculated as the sum of the intrinsic rate of increase (*r*) and death rate (*d*) for uninfected hosts (*b* = *r* + *d*) (Civitello et al. 2013; Shocket et al. 2018a).

Transmission rate (β) was estimated using the numbers of infected and uninfected individuals from each temperature and pathogen treatment using a binomial distribution in a likelihood function to model the number of uninfected hosts in each jar, where the probability of remaining uninfected (*P*) is

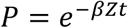

where *Z* is the density of pathogen spores and *t* is the length of the infection period, which allowed us to estimate transmission rate, β (see Shocket et al. 2018a and the supplementary material therein for details of how this likelihood function is derived). In our estimates of transmission rate, individuals were only scored as being infected if they became infected and went on to produce mature transmission spores. Finally, mature spore loads, σ, were quantified for each infected individual as described above.

For all parameters and derived traits we used JAGS (*R2jags* package: Plummer 2003; Su and Yajima 2009) to calculate Bayesian posterior distribution estimates for each trait in turn (as in Shocket et al. 2018a). Two parameters that contribute to our indicator of the potential for disease spread (*R_0_*), host populations carrying capacity, *c*, and spore degradation rate, *m*, were set as constants for all treatments (*c* = 0.01 and *m* = 0.9, taken from Civitello et al. 2013; Shocket et al. 2018a) as they were not measured in this experiment. Whilst it is conceivable that these parameters could vary with temperature exposure, particularly spore degradation (*m*), neither was possible to quantify in these experiments.

Finally, to calculate *R_0_* for each pathogen and temperature treatment, we incorporated the Bayesian posterior estimates of each of the parameters described above into our derived equation for *R_0_*. By incorporating the posterior estimates for each calculated trait in turn, we allowed the propagation of error in our estimates of each trait into our final estimates of the potential for disease spread, *R_0_*.

### Additional statistical analysis

All analyses were conducted in R (v. 3.6.2; R Development Core Team, https://www.R-project.com). Figures were produced using *ggplot2* (Wickham 2016) and *cowplot* (Wilke 2019).

We investigated the effect of infection and thermal acclimation on heat knockdown times by fitting a linear mixed effect model (*nlme* package; Pinheiro et al. 2018). Maternal acclimation temperature (2 levels: 20°C or 25°C), focal acclimation temperature (2 levels: 20°C or 25°C), and pathogen treatment (4 levels: pathogen genotype C1, C14 and C20, or uninfected controls) and their interactions were fitted as fixed effects, while assay run was treated as a random effect. To account for heteroscedasticity in the residual variance in this model, residual variance was allowed to vary independently at the level of the focal acclimation temperature using the *‘VarIdent’* function (*nlme* package: Pinheiro et al. 2018; but see Zuur et al. 2009). The significance of fixed effects were then tested using analysis of variance (ANOVA Type III; *car* package: Fox and Weisberg 2018).

Next, we investigated how various host and pathogen fitness traits varied by both temperature treatment and pathogen exposure. Host lifespan and host total lifetime fecundity were both log transformed and then analysed using ANOVA with maternal temperature, focal temperature, and pathogen treatment (including all higher order interactions) fit as fixed effects. In these analyses of host traits, we included all individuals exposed to a pathogen regardless of whether they ultimately lead to infections with mature transmission spores because here we were focussed on virulence to the host regardless of whether an infection was successful for the pathogen. To analyse within-host pathogen fitness we first predicted the probability that each pathogen genotype would infect and go on to produce mature transmission spores by running a binomial generalized linear model for each treatment combination. Successful infection probability and standard errors were then extracted using the ‘*emmeans*’ function (*emmeans* package: Lenth 2020). We then analysed within-host spore loads for successful infections (those that produced mature transmission spores) using ANOVA with the same fixed effect structure described above. Significance of fixed effects were tested for each linear model using Type III ANOVA (white-corrected to account for residual heteroscedasticity).

For our estimation of disease spread (*R_0_*), we used JAGS to produce posterior distributions for each contributing trait/parameter (see above). Our standard JAGS settings included 75000 iterations, 30000 burn-in, thinning of 16, and 3 individual chains. For each trait we used semi-informative priors, and set the Bayesian posteriors to follow the appropriate distributions. For the Bayesian estimate of transmission rate (β) and the GLM for infection probability, one treatment achieved a 100% infection rate in our experiment (pathogen C20, temperature treatment 20°C −20°C), so to allow more reasonable point estimates and error to be calculated we adjusted this treatment to include one uninfected individual. Due to differences in early survival, handling errors, and male individuals set up unintentionally, sample sizes for the different treatment combinations and disease traits varied between 17 and 26.

## Results

### Disease, acclimation, and thermal tolerance

We first looked at how infection would impact heat knockdown times in combination with both maternal and focal thermal acclimation. We found that the way in which infection mediated host thermal tolerance was highly dependent on prior thermal experience (Figure 1), and determined by a three-way interaction between maternal temperature, focal temperature, and pathogen treatment (Table 1). The clearest effect was that of focal temperature, where individuals directly exposed to 25°C showed a clear improvement in knockdown times across all pathogen treatments compared to individuals exposed to a focal temperature of 20°C (Figure 1). For example, control individuals exposed to a focal temperature of 25°C saw a two-fold increase in knockdown times compared to controls acclimated to 20°C. We also found that pathogen exposure reduced thermal limits compared to controls, but only in the focal 25°C treatments, and that the precise magnitude and direction of this effect depended on the specific pathogen genotype involved (Figure 1). The effect of maternal acclimation prior to infection, contributing to the three-way interaction, was much more subtle, and probably driven by the slight increase in knockdown times for infected individuals in the 20°C-20°C temperature treatment, as well as variation in the reduction in knockdown times between pathogen genotypes at 25°C (Figure 1). There was also a notable increase in the variance in knockdown times across the two focal temperatures, where both control and infected treatments saw far greater variance after exposure to 25°C focal temperature (Figure 1).

**Figure 1:**
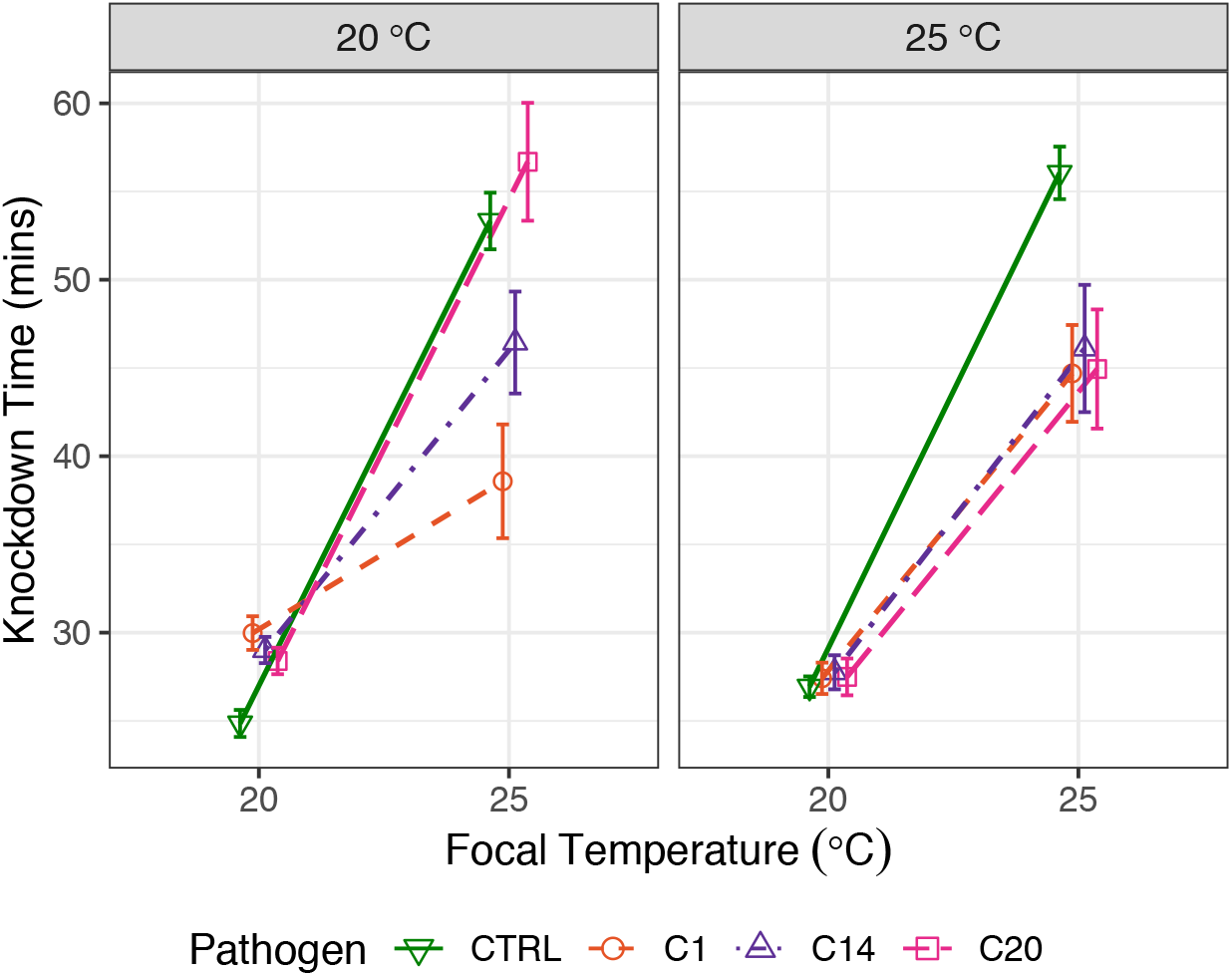
The effect of thermal acclimation on heat knockdown times. Knockdown time was measured for *Daphnia* infected with one of three pathogen genotypes (C1, C14 or C20) or uninfected (CTRL). Each facet represents the maternal/developmental thermal acclimation temperature treatment pre-infection, while the focal temperature was experienced by experimental animals from birth, including over the duration of the infection. Points represent treatment means (± SE).

**Table 1:** The effects of maternal acclimation temperature (20°C or 25°C), focal acclimation temperature (20°C or 25°C), pathogen treatment (Control, C1, C14 or C20) and their interactions on heat knockdown times under 37°C static heat shock.

| Trait | Term | $\chi^2$ | *df* | *p*-value |
| --- | --- | --- | --- | --- |
| Knockdown time | Maternal temp | 0.513 | 1 | 0.474 |
|  | Focal temp | 399.944 | 1 | **< 0.001** |
|  | Pathogen | 14.683 | 3 | **0.002** |
|  | Maternal temp x focal temp | 0.005 | 1 | 0.945 |
|  | Maternal temp x pathogen | 11.090 | 3 | **0.011** |
|  | Focal temp x pathogen | 32.550 | 3 | **< 0.001** |
|  | Maternal temp x focal temp x pathogen | 11.465 | 3 | **0.009** |

### Individual performance: host fitness

Both temperature and pathogen treatments had a significant impact on host lifespan, leading to a three-way interaction between all treatments (Table 2a; Figure 2a). Exposure to the 25°C focal temperature reduced lifespan for both control and exposed individuals compared with the 20°C focal temperature (Figure 2a). However, individuals exposed to a pathogen at 20°C saw a greater relative decrease in lifespan compared to controls than individuals at 25°C (Figure 2a). This led to uninfected controls exposed to 25°C having a similar lifespan to infected individuals at 20°C. Although we found a significant three-way interaction between maternal temperature, focal temperature and pathogen treatment, the effect of maternal temperature was very subtle (Figure 2a).

**Table 2:** The effects of maternal/developmental temperature (20°C or 25°C), focal temperature (20°C or 25°C), pathogen treatment (CTRL, C1, C14 or C20) and all interactions on a) host lifespan, b) host lifetime fecundity, c) pathogen infection success probability, and d) within host pathogen spore loads.

| Trait | Term | *F or $\chi^2$* | *df* | *p*-value |
| --- | --- | --- | --- | --- |
| (a) Host lifespan | Maternal temp | 7.028 | 1 | **0.008** |
|  | Focal temp | 605.015 | 1 | **< 0.001** |
|  | Pathogen | 91.240 | 3 | **< 0.001** |
|  | Maternal temp x focal temp | 0.613 | 1 | 0.434 |
|  | Maternal temp x pathogen | 2.956 | 3 | **0.032** |
|  | Focal temp x pathogen | 4.837 | 3 | **0.003** |
|  | Maternal temp x focal temp x pathogen | 3.160 | 3 | **0.025** |
| (b) Host fecundity | Maternal temp | 0.263 | 1 | 0.608 |
|  | Focal temp | 156.177 | 1 | **< 0.001** |
|  | Pathogen | 729.784 | 3 | **< 0.001** |
|  | Maternal temp x focal temp | 6.324 | 1 | **0.012** |
|  | Maternal temp x pathogen | 3.590 | 3 | **0.014** |
|  | Focal temp x pathogen | 1.241 | 3 | 0.295 |
|  | Maternal temp x focal temp x pathogen | 4.257 | 3 | **0.006** |
| (c) Infection success | Maternal temp | 0.135 | 1 | 0.714 |
|  | Focal temp | 49.42 | 1 | **< 0.001** |
|  | Pathogen | 0.292 | 2 | 0.864 |
|  | Maternal temp x focal temp | 0.530 | 1 | 0.466 |
|  | Maternal temp x pathogen | 3.005 | 2 | 0.223 |
|  | Focal temp x pathogen | 4.573 | 2 | 0.102 |
|  | Maternal temp x focal temp x pathogen | 7.445 | 2 | **0.024** |
| (d) Spore loads | Maternal temp | 3.797 | 1 | 0.053 |
|  | Focal temp | 19.539 | 1 | **< 0.001** |
|  | Pathogen | 10.445 | 2 | **< 0.001** |
|  | Maternal temp x focal temp | 0.913 | 1 | 0.340 |
|  | Maternal temp x pathogen | 0.846 | 2 | 0.431 |
|  | Focal temp x pathogen | 0.958 | 2 | 0.386 |
|  | Maternal temp x focal temp x pathogen | 0.357 | 2 | 0.700 |

**Figure 2:**
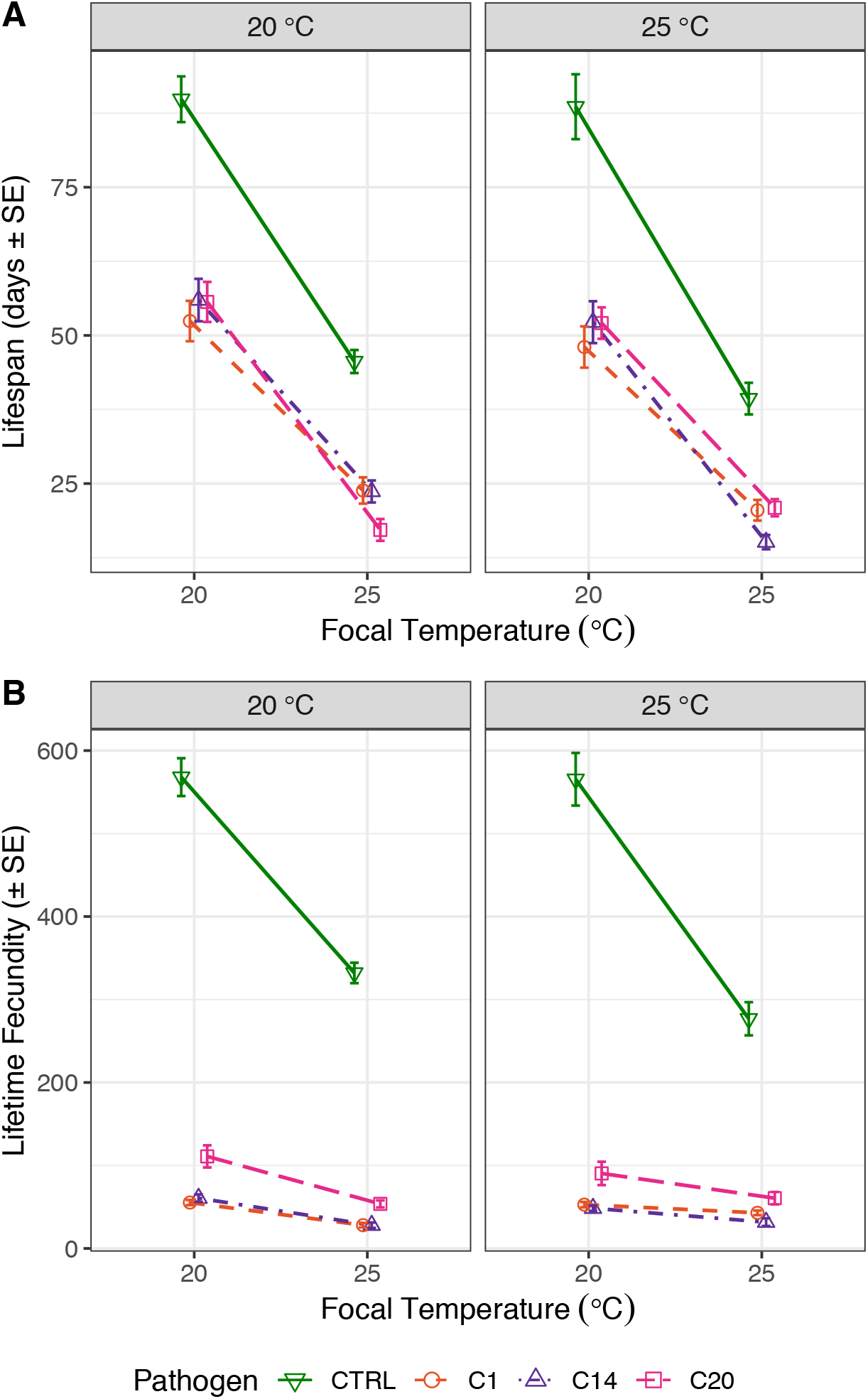
The effect of thermal acclimation on fitness traits of infected and uninfected hosts. A) lifespan, and B) lifetime fecundity, were measured for *Daphnia* infected with one of three pathogen genotypes (C1, C14 or C20) or uninfected (CTRL). Each facet represents the maternal/developmental thermal acclimation temperature treatment pre-infection, while the focal temperature was experienced by experimental animals from birth, including over the duration of the infection. Points represent treatment means (± SE).

Similarly, the interaction between temperature treatments and pathogen exposure had a significant effect on host lifetime fecundity (Table 2b; Figure 2b). Control individuals had greatest fecundity when exposed to a focal 20°C, almost twice the fecundity of control individuals exposed to 25°C, with the greatest reduction occurring for individuals exposed to both maternal and focal 25°C (Figure 2b). Pathogen exposure severely reduced host lifetime fecundity across pathogen genotypes and temperature treatments, with the lowest fecundity seen for individuals exposed to 25°C, although this was only marginally lower than pathogen exposed individuals that were reared at 20°C (Figure 2b). Again the significant contribution of maternal temperature to lifetime fecundity was small, likely driven by a slight increase for control individuals in the maternal 20°C treatment compared to maternal 25°C (Figure 2b).

### Individual performance: within-host pathogen fitness

We next assessed within-host pathogen fitness as both the probability that a pathogen could infect a host and produce mature transmission spores, and also the within host mature spore loads. Successful infection probability was affected by a significant three-way interaction between temperature treatments and pathogen genotype (Table 2c). Successful infection probability was overall greatest for individuals reared at a focal 20°C, with a drop in infection rate for all pathogen genotypes reared at 25°C (Figure 3a). Across maternal temperature treatments, we saw a rank order shift in pathogen genotypes when individuals were subsequently reared at the focal temperature 25°C (Figure 3a). Infection success for two pathogen genotypes dropped as low as 25% at the focal temperature 25°C, but the particular genotype involved depended on the maternal temperature treatment (*i.e*. C20 at 20°C-25°C and C14 at 25°C-25°C; Figure 3a).

**Figure 3:**
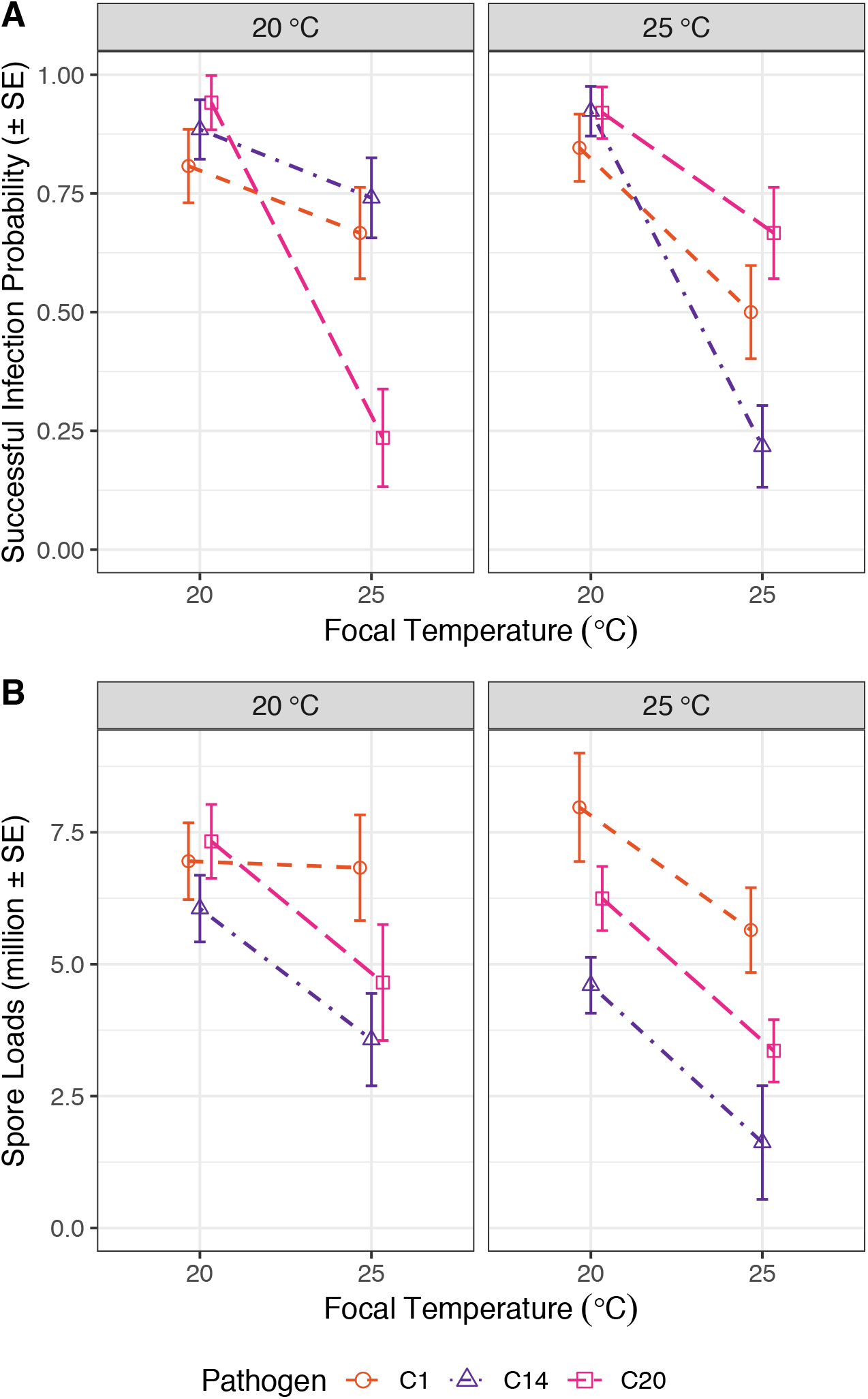
The effect of thermal acclimation on fitness traits of three pathogen genotypes. A) Successful infection probability calculated via a binomial generalized linear model, and B) Within host pathogen spore loads, were measured in hosts infected with one of three pathogen genotypes (C1, C14 or C20). Each facet of the figure represents the maternal and developmental thermal acclimation temperature treatment, while the focal temperature was experienced by experimental animals from birth, including over the duration of the infection. Points represent treatment means (± SE).

Mature spore loads were not driven by any interactions between treatments, and instead were determined by the direct effects of focal temperature and pathogen genotype (Table 2d). Overall we saw a reduction in mature transmission spores in individuals exposed to a focal 25°C, but across temperature treatments each pathogen genotype differed but varied reasonably consistently (Figure 3b), with pathogen genotype C1 generally producing more spores at host death than C14 and C20, respectively.

### Population performance: host population dynamics, pathogen transmission, and disease spread

Using our disease model, we next parameterised a metric of a pathogens potential to spread in a susceptible population (*R_0_*), and investigated the relative contributions of various parameters of host and pathogen population dynamics to disease spread, and how these various parameters were influenced by thermal acclimation. We found that increasing the focal temperature from 20°C to 25°C considerably increased the birth rates of uninfected (susceptible) hosts by around 15%, with similar effects across maternal acclimation temperatures (Figure 4a). Similarly, host death rates also increased with increasing focal acclimation temperature in uninfected individuals, although to a much smaller magnitude than that of birth rates (Figure 4b). However, the contribution of births and deaths of susceptible hosts to our model of disease spread, (b – d)/b, rendered the increase in births at 25°C unimportant, and consequently we saw no difference in the contribution to susceptible host population density across any of our thermal acclimation treatments (Figure 4c).

**Figure 4:**
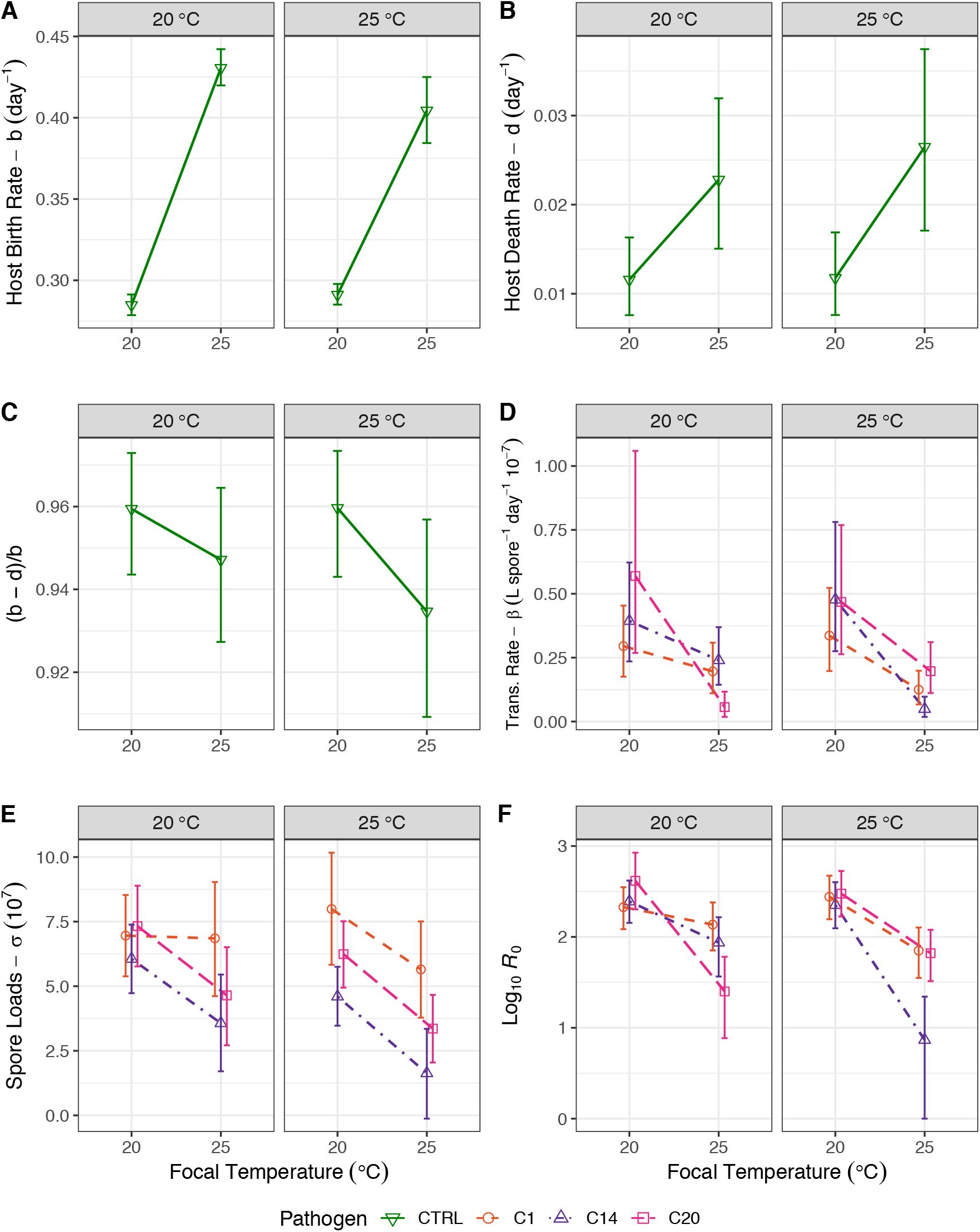
The effect of thermal acclimation on parameters and derived traits from a model of disease spread. A) Birth rate of uninfected hosts (*b*), B) death rates of uninfected hosts (*d*), C) uninfected hosts contribution to disease spread in our model, D) transmission rate (β) for each pathogen genotype, E) spore loads (σ) for each pathogen genotype, F) our composite measure of disease spread (*R_0_*). Shown are the Bayesian estimated posterior means of each parameter or trait for each treatment (with 95% credible intervals) estimated using JAGS. Our metric of disease spread (*R_0_*) is shown on the log10 scale for graphical clarity, and the lower credible interval of pathogen C14 in temperature treatment 25-25 was set to zero (the lowest possible level for this metric) as the raw lower credible interval was negative.

For the epidemiological parameters that were included in our metric of disease spread, pathogen transmission rate was generally higher when hosts were exposed to the focal acclimation temperature of 20°C (Figure 4d), with a reduction in transmission rate at 25°C. We also saw a rank order shift across pathogen genotypes in transmission rate between the two maternal/developmental temperatures when pathogens were subsequently exposed to 25°C (Figure 4d). Our Bayesian posteriors for spore loads (Figure 4e) show the same pattern to our raw data for spore loads (see Figure 3b), where there was an overall decrease in mature spore production at the 25°C focal acclimation temperature, but with similar trends across pathogen genotypes and maternal acclimation temperatures (Figure 4e).

Finally, by combining these parameters, we were able to estimate a metric for the potential of a pathogen genotype to spread in a population (*R_0_*). Like most of our traits, the clearest effect was that of the focal acclimation temperature, where we saw a considerable decrease in *R_0_* at 25°C (Figure 4f). Indeed, for most pathogen genotypes, the potential for disease spread was around an order of magnitude greater at a focal acclimation temperature of 20°C compared to 25°C. This severe reduction in the potential for disease spread at warmer temperatures appears to be driven almost entirely by the effects of temperature on pathogen transmission rate and spore production. We also see a rank order shift in pathogen genotypes across maternal/developmental temperature at the focal acclimation temperature 25°C, which likely represents the effects of maternal acclimation on transmission rate (Figure 4d) influencing the potential of disease spread across maternal acclimation temperatures (Figure 4f).

## Discussion

With both average temperatures and extreme thermal events increasing with global change, will hosts or their pathogens ultimately be the winners or losers? The success of a host or a pathogen population depends on processes that occur at the individual level, including host fecundity and lifespan, and the ability for a pathogen to reproduce within a host, but also on the balance between birth and death rates of a host population and a pathogen’s capacity to spread between susceptible hosts (Anderson and May 1991; Mideo et al. 2008; Hall and Mideo 2018); processes that are properties of the population as a whole. The impact of a pathogen on the thermal limits of its host, however, must also be considered. As extreme temperatures are likely to put greater pressure upon population than changes in average temperatures (Kingsolver and Woods 2016; Sunday et al. 2019), thermal limits, and their modification by infection (Greenspan et al. 2017; Hector et al. 2019), will be vital for population persistence (e.g. Bush et al. 2016). Yet, studies that integrate changes in thermal limits with an exploration of host and pathogen fitness at both the individual and population levels are rare.

In this study, we sought to address how temperature exposure can alter host and pathogen performance by exploring how different types of thermal acclimation could mediate the individual and population level responses of hosts and pathogens to temperature, and comparing this to changes in the thermal limits of both uninfected and infected hosts. We focused on thermal acclimation that occurs during the previous generation and early development, as well as the direct exposure of host and pathogens to warmer temperatures during infection. Our results highlight the importance of considering the impact of thermal acclimation on multiple components of host and pathogen fitness across both the individual to population level scales. Indeed, we find complex and contrasting effects of thermal acclimation on these different measures of performance, particularly for hosts. We discuss our findings in regard to host and pathogen individual performance, the impact of temperature on pathogen virulence, and the population level dynamics of host and pathogens under warmer thermal conditions.

### Thermal acclimation improves thermal tolerances but with a cost to both host and pathogen individual performance

Thermal acclimation to warmer temperatures allows individuals to shift their thermal optima or maxima, potentially acting as a buffer against future thermal stress (Sinclair et al. 2016; Sgrò et al. 2016; Rohr et al. 2018). Here, thermal acclimation as a result of direct exposure to warmer temperatures substantially increased thermal tolerance of uninfected animals. For example, in uninfected individuals, thermal acclimation to warmer temperatures increased knockdown times by up to 30 minutes (Figure 1). This increase in knockdown times is far greater than the variation we see in thermal tolerance across large geographic ranges in *Drosophila melanogaster* (Hoffmann et al. 2002; Sgrò et al. 2010; Lasne et al. 2018; Hector et al. 2020), and is similar to latitudinal variation and the effects of thermal acclimation seen in other *Daphnia* populations (Williams et al. 2012; Yampolsky et al. 2014). In addition, a 30 minute increase in knockdown times after acclimation equals the reduction in knockdown times seen in severely infected *Daphnia* raised at their normal culturing temperatures (Hector et al. 2019). Here, even in infected individuals we saw thermal acclimation enhancing host thermal tolerance, but to a lesser extent than uninfected individuals (an average of 15 minutes increase compared to infected individuals at 20°C). Thermal acclimation, therefore, appears to better prepare both infected and uninfected animals for the pressure of extreme thermal events through plastic shifts in their thermal performance.

For both the host and pathogen, however, the improvement in thermal tolerances with acclimation came with a significant cost to other measures of individual performance. Exposure to 25°C led to substantial reductions in the lifespan and lifetime fecundity of both healthy and infected individuals (Figure 2). Simultaneously, at warmer temperatures, within-host pathogen performance was also low. Both the probability of infection success and within-host spore loads substantially dropped at warmer thermal acclimation temperatures. Indeed, for some pathogen genotypes infection success dropped as low at 25% when infections took place at 25°C, suggesting that at 25°C, while pathogen exposure could have severe impacts on host fitness across most individuals (Figure 2), the proportion of infections that successfully led to mature transmission spores could be very low (Figure 3a). This highlights how thermal acclimation can have opposing impacts on thermal stress resistance and other fitness related traits for both hosts and pathogens (e.g. Cavieres et al. 2020).

### The damage caused by a pathogen is trait-specific and dependent on thermal acclimation

The decline in host fitness that a pathogen causes, known as virulence, is normally assessed in terms of reductions in lifespan or reproduction (Frank 1996; Day 2002; Alizon et al. 2009; Cressler et al. 2016). However, in the context of global change and extreme thermal events, a pathogen’s virulence could equally be extended to include changes in a host’s thermal tolerance. At 20°C we observed that infection resulted in a negligible or even slightly beneficial change in knockdown times relative to uninfected controls, but at 25°C individuals experienced up to a 15-minute reduction in thermal limits as a result of infection. In contrast, however, infected individuals at lower temperatures experienced the greatest virulence in terms of reductions in both lifespan and fecundity relative to uninfected controls. For example, individuals exposed to a pathogen at 20°C experienced a reduction in lifespan of approximately 35 days relative to uninfected individuals, compared to a reduction of around 20 days at 25°C.

By considering the impacts of thermal shifts on the virulence of the pathogen, we see that the damage a pathogen causes its host at warmer temperatures can manifest as a greater reduction in thermal tolerance but a relatively smaller reduction in other, more commonly assessed, fitness traits. These results highlight the complex and contrasting ways in which infection and thermal acclimation can interact to impact various aspects of host performance (Raffel et al. 2013; Manzi et al. 2019; Ferguson and Sinclair 2020). They also highlight a tension between resistance to infection and thermal stress (Hector et al. 2019), such that individuals exposed to 20°C had little to no impact of infection on their thermal tolerance, but overall their thermal tolerance was considerably lower than individuals exposed to 25°C who did nevertheless experience the virulent effect of infection.

### Contrasting host and pathogen performance across individual and population level scales

Our results so far suggest that the improved thermal tolerance when exposed to warmer temperatures comes with a substantial burden to the individual performance of both host and pathogens. Yet, as we show, individual performance metrics can be misleading when population persistence instead depends on vital rates such as growth rates for a host population or the between-host spread of a pathogen (Agha et al. 2018; Shocket et al. 2018a). Important for host population growth is the intrinsic rate of increase, a metric for the age specific fecundity of a population, which can itself be partitioned into the relative contributions of intrinsic birth and death rates to population growth (McCallum 2000; Civitello et al. 2013; Shocket et al. 2018a). In contrast, the success of a pathogen population is dependent on its ability to spread through a host population, encompassed by the parameter *R_0_* (Anderson and May 1986), which depends on both host population dynamics (a combination of birth and death rates) along with key epidemiological traits including pathogen transmission potential and within-host spore proliferation.

The projections of host and pathogen population level performance confirmed how warmer temperatures can accelerate the pace of life for both hosts and pathogens (Angilletta et al. 2004; Fels and Kaltz 2006; Vale et al. 2008; Stoks et al. 2014; Cuco et al. 2018). For the host this led to earlier reproductive output, and higher birth rates as a result (Fig. 4a), but also higher intrinsic death rates (Fig. 4b). The net result however was that these two factors cancel each other out (Fig. 4c). The combination of birth and death rate in our model meant population growth rate of susceptible hosts, ‘(b-d/b)’, was equivalent across all both temperatures, as indicated by overlapping credible intervals in Figure 4c. The shift towards earlier reproductive output and a capacity for faster population growth, therefore, appears to completely offset the severe loss of lifetime fecundity and lifespan that each individual experienced at warmer acclimation temperatures.

In contrast, warmer temperatures reduced pathogen success in terms of *R_0_*, reinforcing the decline in fitness observed for a pathogen within a host. Indeed, the potential for the spread of a pathogen was around an order of magnitude lower at 25°C compared with at 20°C (Fig. 4f). The negligible impact that warmer temperatures had on host vital rates negated any benefit that high birth rates may have provided by enhancing the *R_0_* value of a pathogen. Instead, the decline in pathogen success at the population scale was driven entirely by the reductions in pathogen transmission rate and spore loads at warmer temperatures. One possibility is that under warm temperatures the pace of life became too fast for a pathogen and the vastly shortened lifespan of a host came at a high cost to the pathogen in terms of the time allowed for proliferation (see Vale et al. 2008; Vale and Little 2009; Clerc et al. 2015; Hall and Mideo 2018; Shocket et al. 2019). Alternatively, warmer temperatures may have afforded a host an improved immune response (Elliot et al. 2002; Adamo and Lovett 2011; Ferguson and Sinclair 2020), constraining within-host proliferation, and pathogen success.

### Maternal and developmental acclimation had a marginal impact on most measures of host and pathogen performance

Above we have focussed on the results of direct thermal acclimation, because these were by far the strongest effects. In other species, both maternal temperature effects and the effects of temperature during development are known to influences offspring heat resistance, and other aspects of fitness (discussed in Hoffmann et al. 2012; Beaman et al. 2016). For *Daphnia*, maternal effects have also been found to influence offspring fitness, both in terms of fighting infection and fecundity and lifespan (Mitchell and Read 2005; Hall and Ebert 2012; Garbutt et al. 2014; Michel et al. 2016). These previous results suggest that the combination of maternal and developmental thermal exposure should considerably alter offspring thermal tolerance and fitness, particularly in the face of infection. We found significant effects of maternal temperature across most individual performance traits, although the size of these effects was often small. The effect of maternal temperature was most clear when individuals were subsequently raised at 25°C. For example, the relative impact of pathogen exposure on host thermal tolerance varied across maternal acclimation treatments (Fig 1). At the broader scale, maternal temperature had little impact on host population dynamics (e.g. Fig 4c), but produced its most noticeable impact on pathogen infection success and in turn *R_0_*, where the rank order of pathogen genotypes shifted in response to maternal acclimation temperature (Fig. 4f).

These results suggest that the temperature experienced in the parental generation and during early development may have limited impact on overall host performance, where the temperatures experienced directly during life may swamp any carryover effects. This contrasts with what is seen in terrestrial insects such as *Drosophila* where developmental temperatures can be the most important (Kellermann et al. 2017). Maternal acclimation, however, does have a clear effect on which pathogen genotype was most or least successful. As has also been found when altering the temperature over which infection takes place (Fels and Kaltz 2006; Vale et al. 2008; Vale and Little 2009), variable thermal environments, via maternal or developmental effects, have the potential to maintain genetic variation in pathogen populations by altering infection success and, in turn, the capacity for a pathogen genotype to spread within a population (Garbutt et al. 2014). Overall our results suggest that direct thermal acclimation may set the broad changes in host and pathogen fitness, with the influence of maternal effects coming through more nuanced aspects of a host-pathogen interaction such as rank order shifts in pathogen genotype success.

### Conclusion

In summary, studies of thermal ecology, whether looking at host persistence or disease dynamics, often overlook the capacity for thermal acclimation to act across multiple scales. Here we have shown that warmer temperatures, via thermal acclimation, can benefit a host via increases in thermal tolerance, and any simultaneous costs to individual fitness may be outweighed by an increased capacity for population growth. We also suggest that, for a host, the reduction in thermal tolerance caused by infection at warmer temperatures could be considered as an additional aspect of virulence in light of global change. The outlook for a pathogen under warmer temperatures is, however, bleaker. Within-host pathogen success, and ultimately the potential for disease spread, was severely hampered under warmer temperatures through negative effects on pathogen infection success and spore proliferation. If true for other species, hosts may hold an advantage over pathogens in warmer and more variable environments (but see Shocket et al. 2019). However, to fully understand how shifts in temperature will impact hosts and pathogens requires expanding the study of thermal ecology to include host thermal tolerance and host and pathogen thermal performance, whilst also considering how individual level traits relate to population level performance.

## Acknowledgements

We would like to thank L. Aulsebrook, I. Booksmythe, L. Heffernan, C. Lasne, S. Layh and J. Lush for help with laboratory work. This work was supported by funding from both Monash University and the Australian Research Council.

